# Salidroside and its *in vivo* metabolite tyrosol could act directly on dopamine D2 receptors: a study using RNAseq combined with Connectivity Map analysis

**DOI:** 10.1101/2024.03.03.583234

**Authors:** Ji-Zhou Zhang, Chang Jiang, Jing Han

## Abstract

Salidroside is an active ingredient in traditional Chinese medicine such as Rhodiola. Its *in vivo* metabolite, tyrosol, exist in olive oil and red wine. For a long time, clinical practice and research have shown that both of them have many pharmacological activities, but their targets have not reached unanimous conclusion. The present study proposed that dopamine D2 receptor (DRD2) may be the target of salidroside and tyrosol, by using RNA sequencing combined with Connectivity Map (CMAP) analysis. On this basis, molecular docking, surface plasmon resonance (SPR) and cellular thermal shift assay (CETSA) were used to verify that both salidroside and tyrosol directly bind to DRD2. The results of this study can serve as a guide for further pharmacological research on salidroside and tyrosol.

## 1. Introduction

Salidroside is derived from the traditional Chinese medicine *Rhodiola rosea* L or *Ligustrum lucidum* Ait. Tyrosol, distributed in olive oil and red wine, is aglycone and also metabolite *in vivo* of salidroside. They are both derivatives of phenylethanol. Studies have confirmed that salidroside was extensively metabolized to tyrosol after *i*.*v*. administration immediately, and tyrosol was identified as the main form in all rat tissues, but not salidroside [1]. Both salidroside and tyrosol have abundant *in vivo* activity. Salidroside has wide spectrum of pharmacological activities, like antioxidant, antiinflammatory, anti-cancer, cardio-protective, neuroprotective, anti-depressant, antiaging, anti-diabetic, anti-depressant, anti-hyperlipidemic, immunomodulatory, etc [2]. Tyrosol was also reported many activities: antioxidant, anti-inflammatory, anti-cancer, cardio-protective, neuroprotective, anti-depressant, etc. [3,4]

Despite their numerous physiological and pharmacological functions, as small molecules, the specific targets of salidroside and tyrosol have not yet been confirmed. Therefore, researching and confirming their targets is a breakthrough in studying drug action mechanisms. The CMAP technology, published in Nature in 2007[5], providing an important database in pharmacogenomics research. Based on a large amount of gene expression profile data from human cell lines treated with known target drugs, this online tool based on pattern matching algorithm was developed. It is an emerging technology applied in drug target discovery, drug reposition study and other fields. We attempted to use RNA sequencing to obtain transcriptome tags and compare them with known target drugs on CMAP, in order to speculate on some drugs with the high similarity to salidroside and tyrosol on up- and down-regulating genes, and analyze their target sites.

After first step confirmation of the target, experimental verification is required. In subsequent experiments, we used molecular docking to investigate whether the binding sites of salidroside and tyrosol with the target protein are consistent with known agonists and antagonists. In addition, two experiments, SPR and CETSA, were conducted to confirm the direct binding of salidroside and tyrosol with the predicted target protein.

## 2. Materials and Methods

### 2.1 RNA sequencing

Added 30/60 μM salidroside, or 30/60 μM tyrosol in A549 cells when 50% of the cells were fused, and after 12 hours of treatment, collected samples using Trizol. RNA extraction was performed, and the integrity and total amount of RNA were accurately detected using bioanalyzer (Agilent 2100, Agilent Technologies, Inc., CA, USA) After constructing the library, a preliminary quantification was performed using fluorometer (Qubit2.0, Termo, MA,USA) and the library was diluted to 1.5ng/μl. Subsequently, the insert size of the library was detected using bioanalyzer (Agilent 2100, Agilent Technologies, Inc., CA, USA), and the effective concentration of the library was accurately quantified using qRT-PCR. After passing the test, Illumina (Illumina NovaSeq 6000 Sequencing System, Illumina Inc., CA, USA) sequencing was performed, and 150bp paired-end reads were generated. The basic principle of sequencing was sequencing by synthesis.

Built an index of the reference genome by HISAT2 v2.0.5, and align paired-end clean reads were aligned to the reference genome. Calculated the read counts mapped to each gene using feature Counts (1.5.0-p3) to quantify gene expression levels. Differential expression analysis between two comparison groups were performed by using DESeq2 software (1.20.0). Corrected *P*-value of 0.05 and absolute fold change of 2 were set as the threshold for significantly differential expression.

### 2.2 CMAP

After obtaining the up- and down-regulated gene tags of salidroside and tyrosol on cells compared to the control group, we screened genes that were up- or down-regulated by more than 1.5-fold or 1.2-fold, and imported them into Connectivity Map 2.0. From the obtained drug list, we screened drugs with a score greater than 0.8 or less than −0.8 in Connectivity Map 2.0, which indicated salidroside or tyrosol has high similarity or dissimilarity with these drugs, for further analysis.

### 2.3 Molecular Docking

AutodockTools were used for molecular docking. The molecular structures of ligands (dopamine, salidroside, tyrosol, risperidone) were downloaded from PubChem’s database. The protein structure of the receptor (DRD2) was obtained from paper [6].

The grid energy map was drawn to perform molecular docking. The three-dimensional structure of the ligand-receptor binding and its binding ability vina score were explored.

### 2.4 SPR

Nicoya platform (Nicoya OperSPR Instrument, Nicoya, Kitchener-Waterloo, Canada) was used to performing SPR. Install the COOH chip according to the OpenSPRTM instrument standard operating procedure. Start running at maximum flow rate (150 µL/min) with PBS (pH 7.4) as the detection buffer. After reaching the signal baseline, loaded 200 µL of isopropanol and ran for 10 seconds to remove bubbles. After reaching the baseline, rinsed the sample ring with buffer and emptied it with air. After the signal reached the baseline, adjusted the buffer flow rate to 20 µL/min. Activated the chip with a sample of EDC/NHS (1:1) solution.

Loaded 200µL of DRD2 (Abnova H00001813-Q01, Abnova, Taipei, China) for 4 minutes, washed the sample loop with buffer, and emptied it with air. Loaded 200µL of blocking solution, washed the sample loop with buffer (PBS), and emptied it with air. Observed the baseline for 5 minutes to ensure stability. Dopamine, salidroside, and tyrosol are diluted in buffer (PBS) using a gradient, with dopamine gradient of 0.4-0.8-1.6-3.2 mM, salidroside and tyrosol gradient of 0.8-1.6-3.2-6.4 mM, loaded at 20 µL/min, and protein-ligand binding time was 240 seconds, natural dissociation was 260 seconds.

The experimental results were analyzed using TraceDrawer (Ridgeview Instruments ab, Sweden) using the One To One analysis model. The concentration gradient binding curves of the three test substances with DRD2 were plotted, and the following kinetic and affinity parameters were calculated: binding rate constant, dissociation rate constant, and dissociation equilibrium constant (affinity constant).

### 2.5 CETSA

A549 cells were seeded into a six-well plate, and treated 120μM salidroside or tyrosol for 0.5h. 300μl cold PBS was added to each well, and the adherent cells were directly scraped with a spatula. The cell suspension was collected in Eppendorf tubes. The cell suspension was heated using a PCR instrument (LongGene T10S, LongGene, Hangzhou, China) with a temperature gradient of 45°C-50°C-55°C-60°C-65°C-70°C-75°C for 3min. The cell suspension was repeatedly frozen and thawed five times in liquid nitrogen/room temperature, in order to fully lyse the cells. Centrifuged at 12000×g for 15min, and the supernatant was aspirated, loading buffer was added at a volume of 1/4 for subsequent Western blot detection.

Western blot steps: load 50μl supernatant sample per lane, 180V constant voltage electrophoresis using 7.5% SDS-PAGE, 0.35mA constant current electro-transferred to PVDF membrane in ice bath. The membrane was blocked by 5% skim milk in TBST for 2h. Incubated with DRD2 primary antibody (1:1000, Abcam ab30743, Abcam, Cambridge, UK) overnight at 4°C, then washed in 1×TBST 3 times for 10 min each time, incubated with secondary antibody (1:2000, Boster BA1006, Boster, Wuhan, China) at room temperature for 2h. Washed membrane in 1×TBST 3 times for 10 min each time, then incubated the ECL reagent, photographed, and quantified with ImageJ (LOCI, University of Wisconsin, USA), to draw a temperature-protein relative expression line graph.

## 3. Results

### 3.1 RNA sequencing

We obtained the protein coding genes that were up- or down-regulated by salidroside and tyrosol by more than 1.5 times/1.2 times compared to the Control group after treating cells. The number of genes is shown in Table 1 and Table 2.

**Table 1:** Genes up- or down-regulated more than 1.5 times after treated.

|  | Salidroside<br>60μM | Salidroside<br>30μM | Tyrosol<br>60μM | Tyrosol<br>30μM |
| --- | --- | --- | --- | --- |
| Up-regulated gene numbers | 31 | 15 | 19 | 43 |
| Down-regulated gene numbers | 40 | 33 | 43 | 33 |

**Table 2:** Genes up- or down-regulated more than 1.2 times after treated.

|  |  | **Salidroside**<br>**60μM** | **Salidroside**<br>**30μM** | **Tyrosol**<br>**60μM** | **Tyrosol**<br>**30μM** |
| --- | --- | --- | --- | --- | --- |
| Up-regulated<br>gene numbers |  | 356 | 66 | 78 | 449 |
| Down-regulated<br>gene numbers |  | 293 | 93 | 103 | 85 |

### 3.2 CMAP

We input the up-/down-regulated genes into Connectivity Map 2.0, and summarized several sets of the most representative results. All these screened drugs have effects on DRD2.

Table 3 showed the summarized results of CMAP. A score of 1 indicated a similarity of 100%, and a score of −1 indicated a similarity of −100% (i.e. 100% dissimilarity). Only drugs with a score higher than 0.8 or less than −0.8 are displayed. The target information for all drugs was obtained from the DrugBank website (https://go.drugbank.com). Only those Pharmacoaction marked as “Yes” on the website were selected (meant the functions have been confirmed by previous researchers).

**Table 3:**
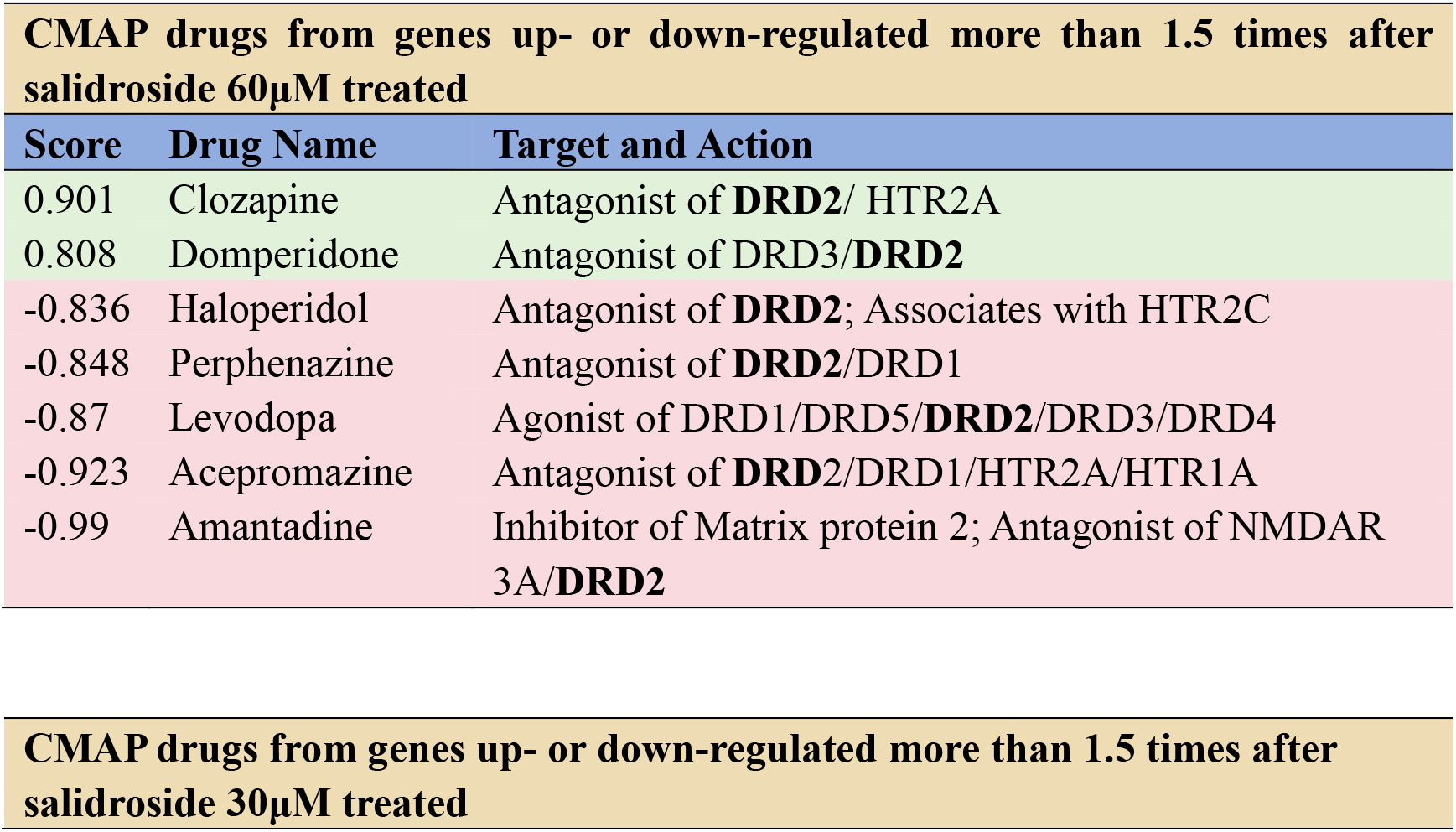
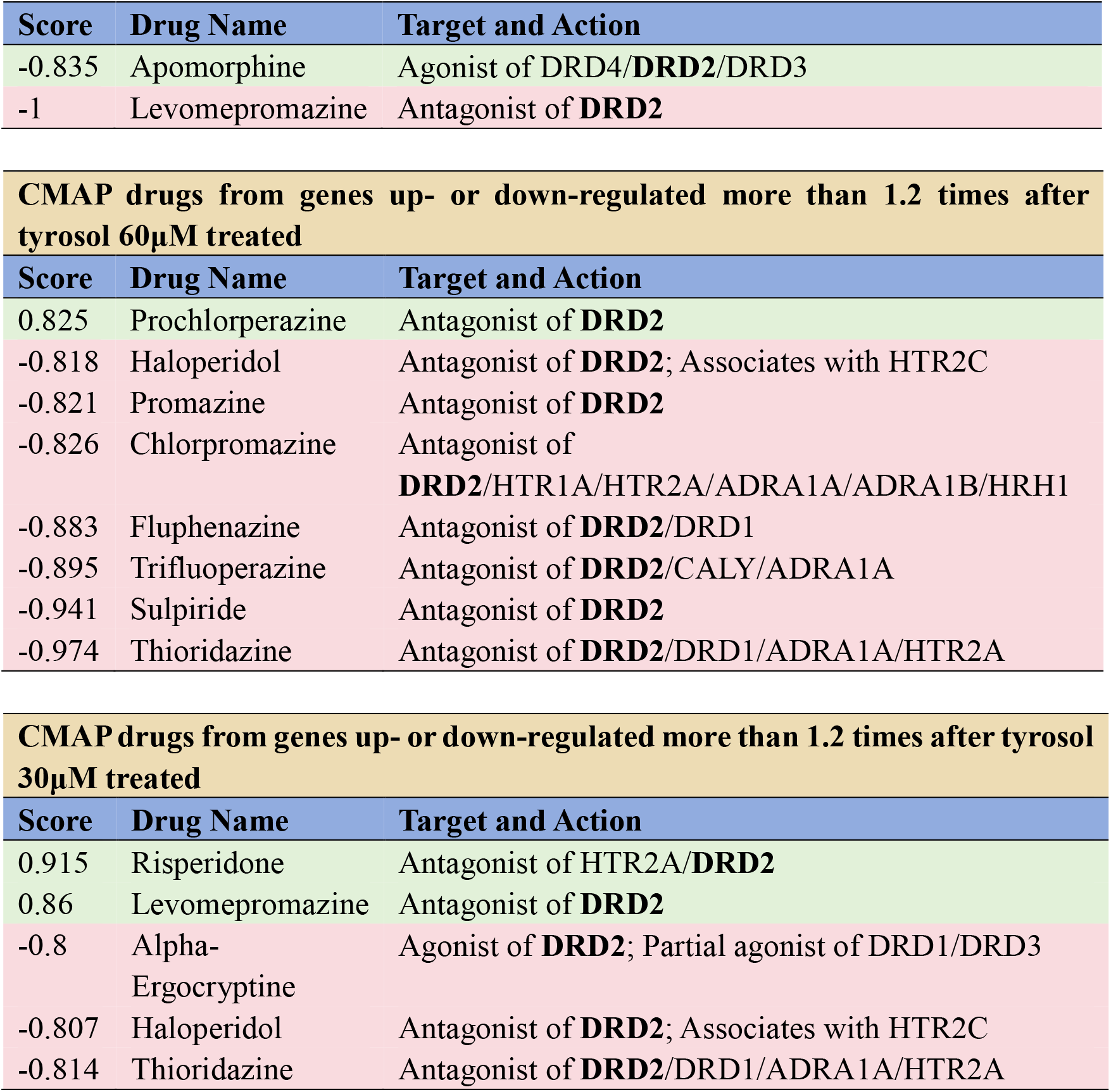
CMAP result from genes up- or down-regulated by salidroside and tyrosol.

The overall results including other drugs are in the appendix.

### 3.3 Molecular docking

Molecular docking results (Figure 1 & Table 4) indicated that both salidroside and tyrosol, as well as dopamine and DRD2 antagonist risperidone, can enter the seven transmembrane domains of DRD2 molecule, known as the “drug binding pocket”. The vina scores obtained from molecular docking are shown in the table. Lower vina scores represented higher binding ability. It can be seen that the vina scores of salidroside and typosol are similar to the score of dopamine.

**Table 4:** The vina scores, the centers and sizes of the docking box of molecular docking with DRD2.

| Molecular | Vina Score | Center |  |  | Size |  |  |
| --- | --- | --- | --- | --- | --- | --- | --- |
|  |  | x | y | z | x | y | z |
| Dopamine | -6.6 | 10 | 4 | -10 | 18 | 18 | 18 |
| Tyrosol | -6.3 | 10 | 4 | -10 | 17 | 17 | 17 |
| Salidroside | -7.6 | 10 | 4 | -10 | 21 | 21 | 21 |
| Risperidone | -12.1 | 10 | 4 | -10 | 26 | 26 | 26 |

**Figure 1.**
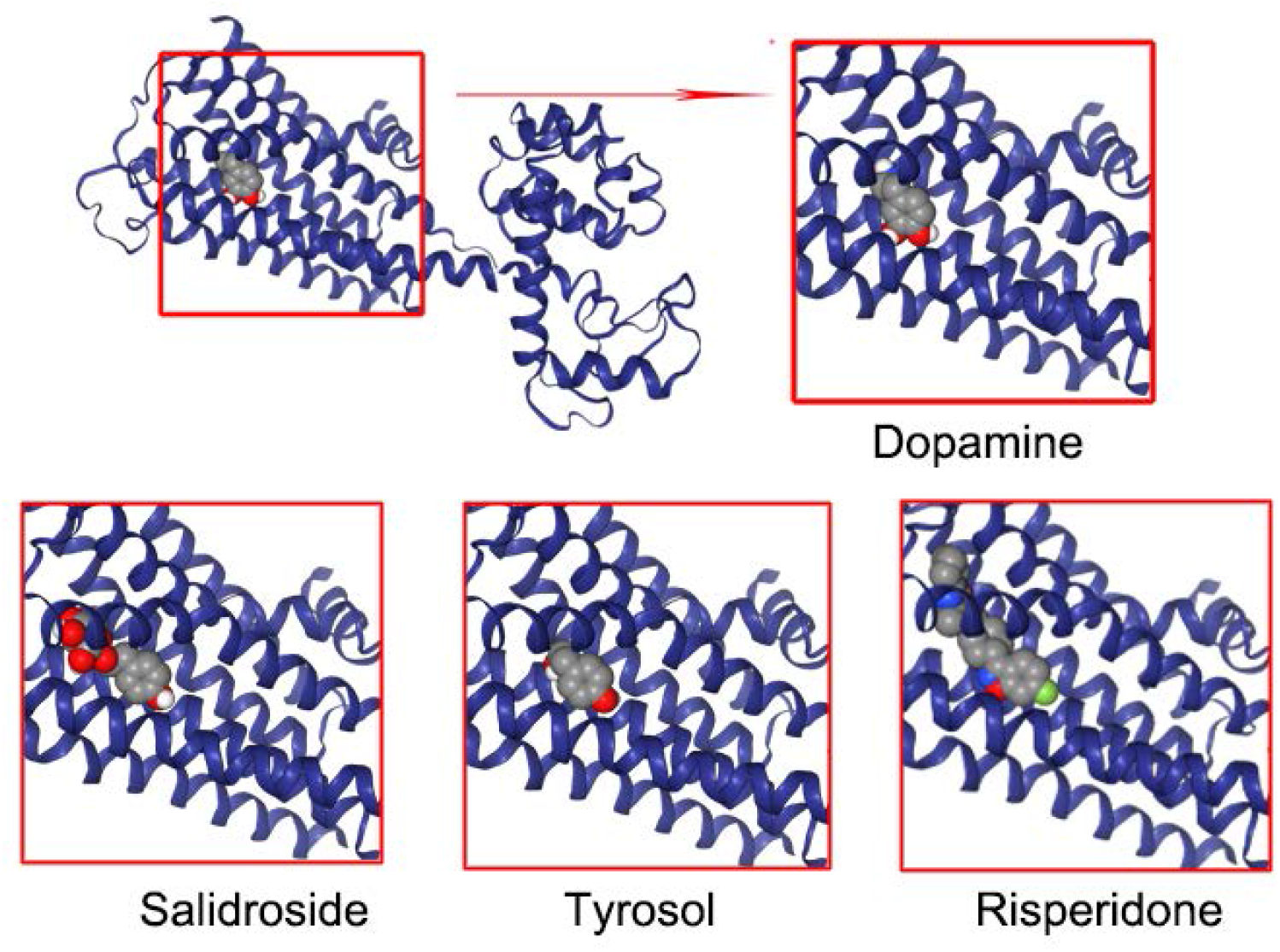
3D diagrams of molecular docking of dopamine, salidroside, tyrosol, risperidone with DRD2. “Drug binding pocket” was enlarged in each diagram.

### 3.4 SPR

Figure 2 shows the concentration-gradient binding curves of dopamine, salidroside, and tyrosol with DRD2. The kinetic and affinity parameters obtained from the curves are shown in Table 5. From the affinity parameters, it can be seen that affinity constant of salidroside and tyrosol for DRD2 are approximately twice that of dopamine.

**Table 5:**
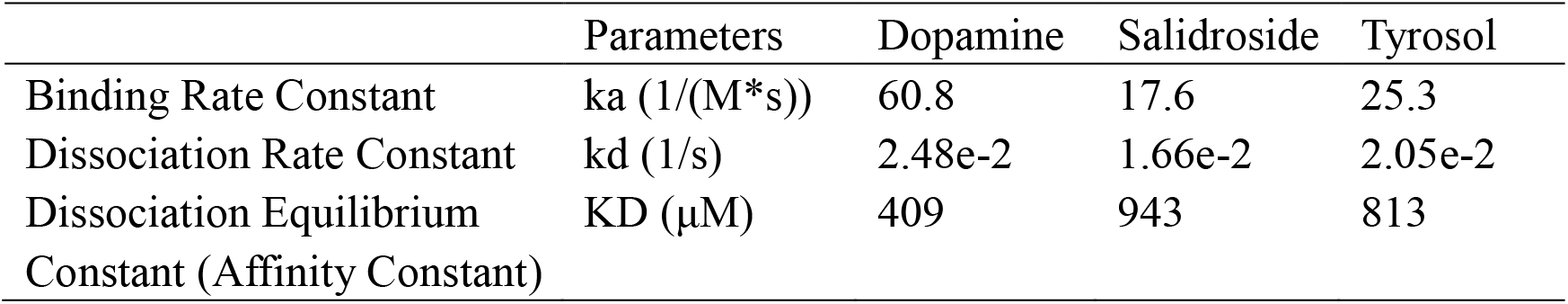
Kinetic and affinity parameters of dopamine, salidroside and tyrosol with DRD2.

|  | Parameters | Dopamine | Salidroside | Tyrosol |
| --- | --- | --- | --- | --- |
| Binding Rate Constant | $k_a$ (1/(M*s)) | 60.8 | 17.6 | 25.3 |
| Dissociation Rate Constant | $k_d$ (1/s) | 2.48e-2 | 1.66e-2 | 2.05e-2 |
| Dissociation Equilibrium Constant (Affinity Constant) | KD (μM) | 409 | 943 | 813 |

**Figure 2.**
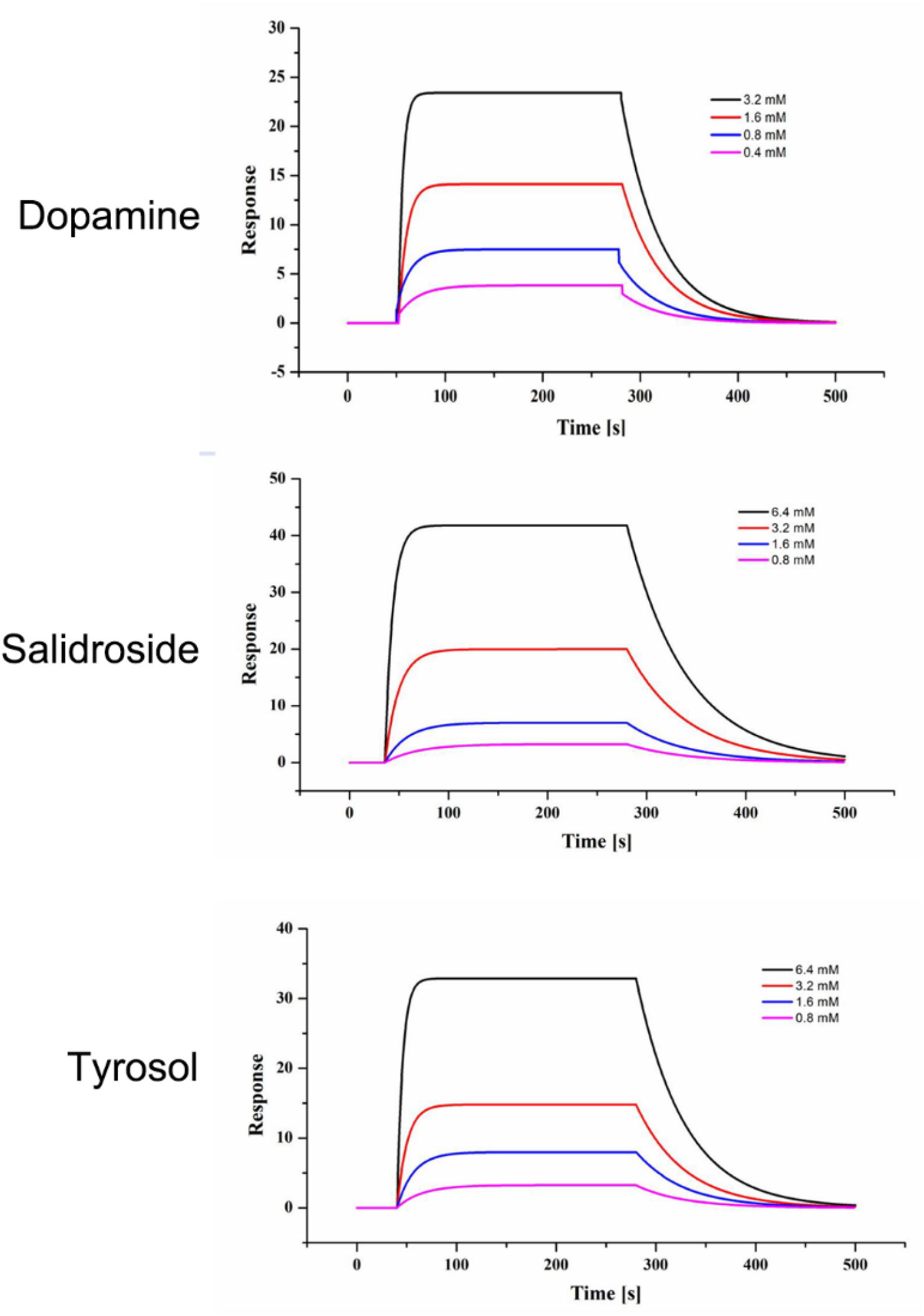
the concentration-gradient binding curves of dopamine, salidroside, and tyrosol with DRD2.

### 3.5 CETSA

As the temperature rose, the DRD2 protein in the cell suspension underwent thermal denaturation, resulting in a decrease in solubility. After centrifugation, the DRD2 content remaining in the supernatant decreased. When DRD2 bound to its ligand, its thermal stability increased, and the rate of decrease in content with temperature slowed down. From Figure 3, it can be seen that compared to the vehicle group, the DRD2 protein content in the supernatant is significant higher after treatment with salidroside or tyrosol 120μM for 0.5h at 50-60°C.

**Figure 3.**
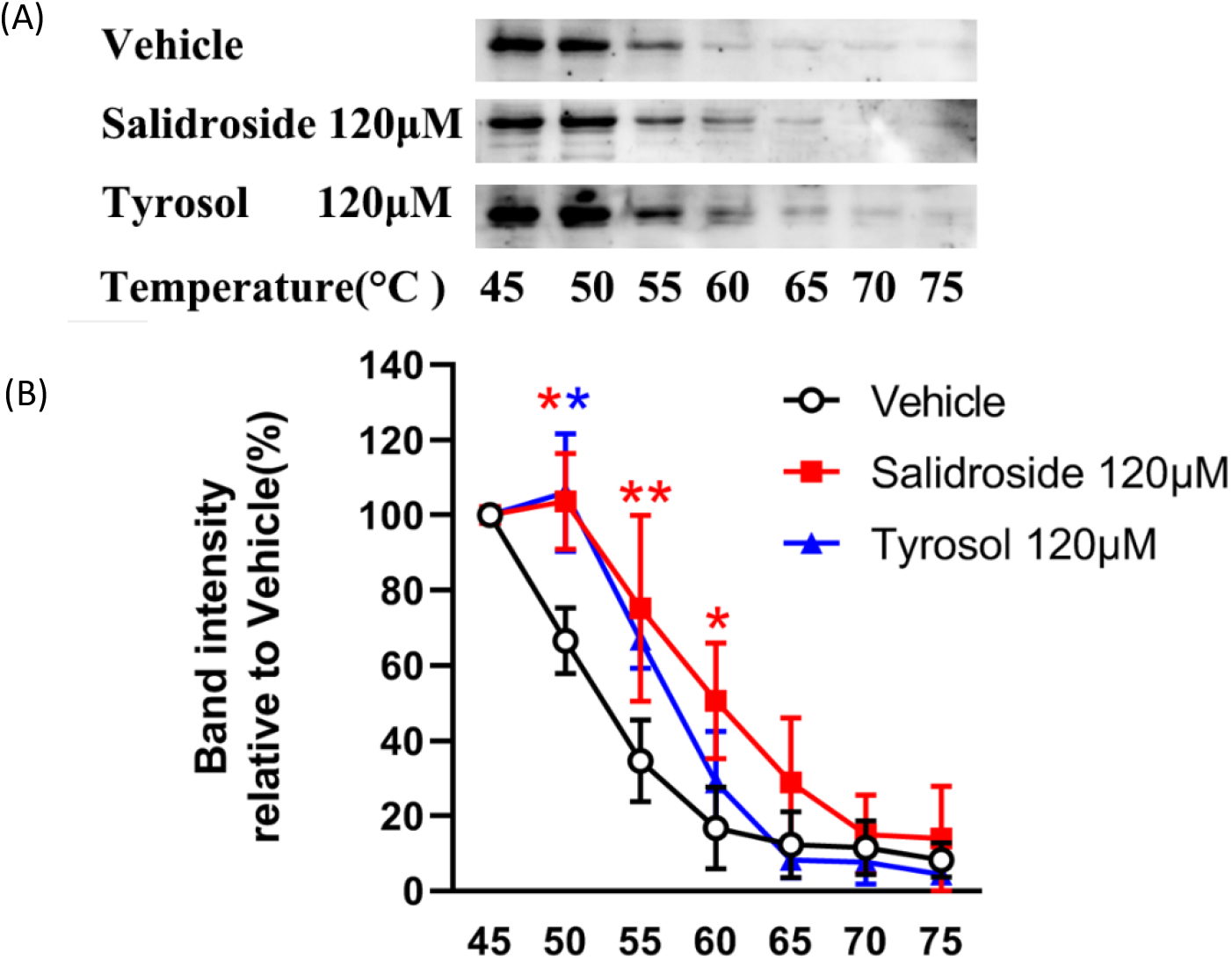
After bound with salidroside or tyrosol, thermal stability of DRD2 protein increased. (A) Western blot analysis of DRD2. (B) DRD2 protein level was quantified. Results were presented as means ± SE. ^*^*P* < 0.05 and ^**^*P* < 0.01, salidroside or tyrosol group compared with the vehicle group.

## 4. Discussion

Through RNA sequencing combined with CMAP analysis, we found the genes up- and down-regulated by salidroside and tyrosol after treated cells similar to some DRD2 related drugs. Therefore, we speculated that DRD2 may be the target of salidroside and tyrosol. Then we used molecular docking to analyze the binding region of salidroside and tyrosol with DRD2, which was consistent with dopamine and DRD2 antagonist risperidone. Next, two experiments at the molecular level of SPR and cellular level of CETSA confirmed that salidroside and tyrosol can directly bind to DRD2, at the molecular and cellular level separately.

The effect of salidroside and tyrosol on DRD2 may be the reason they protect dopaminergic neurons, anti-Parkinson, and mesenchymal stem cells to differentiate into dopaminergic neurons [7–10]. Moreover, this regulation is very rapid. In our published study, we investigated the regulation of salidroside on dopaminergic nervous system after cerebral ischemia. After intraperitoneal injection of salidroside in middle cerebral artery occlusion model rats, the dopamine content in striatum significantly increased, and the contents of its metabolites 3,4-dihydroxyphenylacetic acid and homovanillic acid also increased compared with the model groups, significant differences were observed only 40 minutes later [11]. This is the reason we selected DRD2 as the further research object after preliminary screening in CMAP. Moreover, the structure of tyrosol is similar to dopamine, structural similarity may be the basis for consistent target. Tyrosol is metabolized by monoamine oxidase to 3,4-dihydroxyphenylacetaldehyde, or by aldehyde dehydrogenase to 3,4-dihydroxyphenylacetic acid, and then metabolized by catechol-*O*-methyl transferase to homovanillic acid; similarly, tyramine can be metabolized to tyrosol by monoamine oxidase and aldehyde/aldose reductase [12].

There are multiple advantages of CMAP applied in traditional Chinese medicine. One of them is repositioning traditional Chinese medicine by systematically comparing drug-related gene expression profiles [13]. This function is very useful for researching traditional Chinese medicine, as they are drugs that have been used for centuries and often have complex compositions, and their specific pharmacological properties have not been thoroughly studied. This is fundamentally different from drugs which structure determines properties paradigm. Therefore, CMAP provides a good reference database that allows us to analyze the possible targets of one or more small molecules, thus making the research more clear.

However, CMAP can only provide preliminary research guidance. The results we have obtained so far still cannot determine the direction of regulation of salidroside and tyrosol on DRD2. It is still unclear whether they are agonists or antagonists of DRD2. Which genes are directly related to the regulation of DRD2 are still unknown, because CMAP did not mention these, and only gave answers based on an automated comparison database. Also, the existing results did not indicate enough differences in the regulatory mechanisms of salidroside and tyrosol. Therefore, more cell and animal experiments need to be conducted to further understand how salidroside and tyrosol regulate DRD2, the strength of their effects, and their physiological and pharmacological effects. Additionally, DRD2 may not be the only target of salidroside and tyrosol. These two moleculars may still regulate other targets, which also needs further verification.

## 5. Conclusion

In this study, we used RNA sequencing combined with CMAP to speculate that the target of salidroside and tyrosol is DRD2, and verified the direct binding of salidroside and tyrosol to DRD2 using molecular docking, SPR, and CETSA. The results obtained will provide direction for future research and represent a new approach for the study of single compounds in traditional Chinese medicine or other natural compounds.

## Supporting information

The overall results including other drugs are in the appendix.

## Abbreviations

ADRA: Adrenergic alpha receptor
CETSA: Cellular thermal shift assay
CMAP: Connectivity Map
DR: Dopamine receptor
HR: Histamine receptor
HTR: 5-Hydroxytryptamine receptor
NMDAR: N-methyl-D-aspartate-receptor
PBS: Phosphate buffer
RNA: Ribonucleic acid
SPR: Surface plasmon resonance
TBST: Tris-buffered saline with Tween 20

## Data Availability

The data in the manuscript are from experiment results. More detailed supporting data can be provided upon request to the corresponding author.

## Authors’ Contributions

Jing Han conceived and designed the study. Ji-Zhou Zhang and Jing Han performed the experiments. Chang Jiang performed the molecular docking. Ji-Zhou Zhang and Jing Han wrote the manuscript. All authors read and approved the final content of the manuscript.

## Acknowledgments

This work was supported by grants from Natural Science Foundation of Fujian Province of China (grant no. 2021J01910).

