## Supplementary material for "Salidroside and its *in vivo* metabolite tyrosol could act directly on dopamine D2 receptors: a study using RNAseq combined with Connectivity Map analysis": The overall results including other drugs are in the appendix.

**CMAP drugs from genes up- or down-regulated more than 1.5 times after salidroside 60µM treated**

| Rank | CMAP Name | Dose | Cell | Score |
|---|---|---|---|---|
| 1 | Mefloquine | 10 µM | MCF7 | 1 |
| 2 | Seneciphylline | 12 µM | HL60 | 1 |
| 3 | Securinine | 18 µM | MCF7 | 0.984 |
| 4 | Disulfiram | 13 µM | MCF7 | 0.956 |
| 5 | Pioglitazone | 10 µM | PC3 | 0.937 |
| 6 | Iobenguane | 11 µM | MCF7 | 0.926 |
| 7 | 15-Delta Prostaglandin J2 | 10 µM | MCF7 | 0.919 |
| 8 | Dextromethorphan | 11 µM | MCF7 | 0.918 |
| 9 | Securinine | 18 µM | MCF7 | 0.909 |
| **10** | **Clozapine** | **12 µM** | **HL60** | **0.901** |
| 11 | Dorzolamide | 11 µM | HL60 | 0.898 |
| 12 | Parthenolide | 16 µM | MCF7 | 0.894 |
| 13 | Valproic Acid | 1 mM | HL60 | 0.887 |
| 14 | 15-Delta Prostaglandin J2 | 10 µM | MCF7 | 0.886 |
| 15 | Nocodazole | 13 µM | PC3 | 0.884 |
| 16 | Monorden | 100 nM | MCF7 | 0.881 |
| 17 | Loperamide | 8 µM | PC3 | 0.88 |
| 18 | Natamycin | 6 µM | MCF7 | 0.875 |
| 19 | Flecainide | 8 µM | MCF7 | 0.868 |
| **20** | **Phenelzine** | **17 µM** | **PC3** | **0.863** |
| 21 | Parthenolide | 16 µM | PC3 | 0.863 |
| 22 | Tanespimycin | 1 µM | HL60 | 0.861 |
| 23 | Prestwick-642 | 14 µM | PC3 | 0.858 |
| 24 | Mifepristone | 9 µM | MCF7 | 0.852 |
| 25 | Dydrogesterone | 13 µM | HL60 | 0.849 |
| 26 | Geldanamycin | 1 µM | PC3 | 0.846 |
| 27 | Ciclacillin | 12 µM | MCF7 | 0.843 |
| 28 | Phenoxybenzamine | 12 µM | MCF7 | 0.841 |
| 29 | Dexibuprofen | 19 µM | HL60 | 0.841 |
| 30 | Withaferin A | 1 µM | MCF7 | 0.836 |
| 31 | Disulfiram | 13 µM | PC3 | 0.835 |
| 32 | Prestwick-864 | 35 µM | PC3 | 0.833 |
| 33 | Carcinine | 22 µM | MCF7 | 0.831 |
| 34 | Pregnenolone | 13 µM | HL60 | 0.83 |
| 35 | Trifluridine | 14 µM | MCF7 | 0.83 |
| 36 | Corticosterone | 12 µM | MCF7 | 0.829 |
| 37 | Clioquinol | 13 µM | MCF7 | 0.827 |
| 38 | Withaferin A | 1 µM | MCF7 | 0.827 |
| 39 | Dihydrostreptomycin | 3 µM | HL60 | 0.825 |

| | | | | |
|---|---|---|---|---|
| 40 | Valproic Acid | 1 mM | HL60 | 0.82 |
| 41 | Proguanil | 14 µM | MCF7 | 0.818 |
| 42 | Novobiocin | 6 µM | HL60 | 0.818 |
| 43 | Sulfadimidine | 13 µM | MCF7 | 0.813 |
| 44 | Monorden | 100 nM | MCF7 | 0.812 |
| 45 | Carmustine | 100 µM | MCF7 | 0.812 |
| **46** | **Domperidone** | **7 µM** | **HL60** | **0.808** |
| 47 | Diazoxide | 17 µM | PC3 | 0.807 |
| 48 | 15-Delta Prostaglandin J2 | 10 µM | HL60 | 0.804 |
| 49 | Naringenin | 15 µM | HL60 | 0.804 |
| 50 | Monorden | 100 nM | HL60 | 0.804 |
| 51 | Ritodrine | 12 µM | HL60 | 0.804 |
| 52 | F0447-0125 | 10 µM | MCF7 | 0.804 |
| 53 | Oxaprozin | 14 µM | PC3 | 0.801 |
| 54 | Metolazone | 11 µM | MCF7 | 0.801 |
| 6050 | Calcium Pantothenate | 8 µM | HL60 | -0.804 |
| 6051 | Nitrofurantoin | 17 µM | HL60 | -0.807 |
| 6052 | Pilocarpine | 15 µM | HL60 | -0.808 |
| 6053 | Aminophenazone | 17 µM | HL60 | -0.809 |
| 6054 | Lobelanidine | 11 µM | MCF7 | -0.812 |
| 6055 | Azathioprine | 14 µM | HL60 | -0.814 |
| 6056 | Procainamide | 15 µM | MCF7 | -0.816 |
| 6057 | Neomycin | 4 µM | MCF7 | -0.818 |
| 6058 | Acetazolamide | 18 µM | HL60 | -0.821 |
| 6059 | LY-294002 | 10 µM | HL60 | -0.823 |
| 6060 | Novobiocin | 6 µM | MCF7 | -0.824 |
| 6061 | PHA-00851261E | 10 µM | PC3 | -0.825 |
| 6062 | Isoniazid | 29 µM | PC3 | -0.825 |
| 6063 | Nitrendipine | 11 µM | MCF7 | -0.826 |
| 6064 | Fenbufen | 16 µM | MCF7 | -0.827 |
| 6065 | 0173570-0000 | 10 µM | PC3 | -0.827 |
| 6066 | Vincamine | 11 µM | PC3 | -0.828 |
| 6067 | Dacarbazine | 22 µM | HL60 | -0.831 |
| 6068 | BCB000040 | 10 µM | PC3 | -0.832 |
| 6069 | AG-013608 | 10 µM | MCF7 | -0.834 |
| 6070 | Stachydrine | 22 µM | HL60 | -0.835 |
| **6071** | **Haloperidol** | **10 µM** | **HL60** | **-0.836** |
| 6072 | Oleandomycin | 5 µM | HL60 | -0.836 |
| 6073 | Zalcitabine | 19 µM | HL60 | -0.838 |
| 6074 | CP-645525-01 | 10 µM | PC3 | -0.839 |
| 6075 | Xylazine | 18 µM | PC3 | -0.84 |
| 6076 | Cefalexin | 11 µM | HL60 | -0.841 |
| 6077 | Metrifonate | 16 µM | HL60 | -0.845 |
| 6078 | Perphenazine | 10 µM | HL60 | -0.848 |

| | | | | |
|---|---|---|---|---|
| 6079 | Zimeldine | 10 µM | HL60 | -0.848 |
| 6080 | Tracazolate | 12 µM | MCF7 | -0.851 |
| 6081 | PNU-0230031 | 10 µM | PC3 | -0.851 |
| 6082 | Ketoconazole | 8 µM | HL60 | -0.852 |
| 6083 | Prestwick-665 | 12 µM | PC3 | -0.853 |
| 6084 | Prednisolone | 11 µM | HL60 | -0.858 |
| 6085 | Methoxamine | 16 µM | HL60 | -0.859 |
| 6086 | Iproniazid | 14 µM | MCF7 | -0.87 |
| **6087** | **Levodopa** | **20 µM** | **HL60** | **-0.87** |
| 6088 | Nilutamide | 13 µM | MCF7 | -0.879 |
| 6089 | Levobunolol | 12 µM | HL60 | -0.885 |
| 6090 | Tribenoside | 8 µM | PC3 | -0.886 |
| 6091 | Moxonidine | 17 µM | HL60 | -0.892 |
| 6092 | Ethosuximide | 28 µM | HL60 | -0.896 |
| 6093 | Terbutaline | 7 µM | PC3 | -0.914 |
| **6094** | **Acepromazine** | **9 µM** | **HL60** | **-0.923** |
| 6095 | Prestwick-857 | 12 µM | MCF7 | -0.93 |
| 6096 | Phensuximide | 21 µM | MCF7 | -0.943 |
| 6097 | Propranolol | 14 µM | PC3 | -0.96 |
| 6098 | Metamizole Sodium | 12 µM | MCF7 | -0.972 |
| **6099** | **Amantadine** | **10 µM** | **PC3** | **-0.99** |
| 6100 | Bethanechol | 20 µM | HL60 | -1 |

**CMAP drugs from genes up- or down-regulated more than 1.5 times after salidroside 30µM treated**

| Rank | CMAP Name | Dose | Cell | Score |
|---|---|---|---|---|
| 1 | Dimethadione | 31 µM | PC3 | 1 |
| 2 | Nordihydroguaiaretic Acid | 1 µM | ssMCF7 | 0.861 |
| 3 | Ethosuximide | 28 µM | PC3 | 0.849 |
| 4 | Lobeline | 11 µM | PC3 | 0.846 |
| 5 | Fenbufen | 16 µM | PC3 | 0.801 |
| 6084 | Trichostatin A | 100 nM | MCF7 | -0.803 |
| 6085 | Trichostatin A | 100 nM | MCF7 | -0.809 |
| 6086 | Guaifenesin | 20 µM | PC3 | -0.81 |
| **6087** | **Apomorphine** | **6 µM** | **PC3** | **-0.835** |
| 6088 | Tanespimycin | 1 µM | HL60 | -0.839 |
| 6089 | CP-944629 | 10 µM | PC3 | -0.839 |
| 6090 | Nystatin | 4 µM | PC3 | -0.839 |
| **6091** | **Minaprine** | **11 µM** | **HL60** | **-0.845** |
| 6092 | Prestwick-682 | 6 µM | MCF7 | -0.847 |
| 6093 | Pha-00745360 | 10 µM | MCF7 | -0.854 |
| 6094 | Nifedipine | 12 µM | MCF7 | -0.859 |
| 6095 | Tremorine | 15 µM | PC3 | -0.864 |
| 6096 | CAY-10397 | 10 µM | PC3 | -0.872 |
| 6097 | Hexetidine | 12 µM | PC3 | -0.878 |
| 6098 | Meteneprost | 10 µM | PC3 | -0.88 |
| 6099 | CP-320650-01 | 1 µM | PC3 | -0.934 |
| **6100** | **Levomepromazine** | **9 µM** | **PC3** | **-1** |

**CMAP drugs from genes up- or down-regulated more than 1.5 times after tyrosol 60μM treated**

| Rank | CMAP Name | Dose | Cell | Score |
|---|---|---|---|---|
| 1 | Levopropoxyphene | 7 μM | HL60 | 1 |
| 2 | Metronidazole | 23 μM | PC3 | 0.931 |
| 3 | Merbromin | 5 μM | HL60 | 0.928 |
| 4 | Diethylstilbestrol | 15 μM | HL60 | 0.901 |
| 5 | Estradiol | 100 nM | MCF7 | 0.9 |
| 6 | Cyclic Adenosine Monophosphate | 12 μM | HL60 | 0.895 |
| 7 | Nadolol | 13 μM | HL60 | 0.862 |
| 8 | Benzathine Benzylpenicillin | 4 μM | PC3 | 0.845 |
| 9 | Estradiol | 15 μM | HL60 | 0.845 |
| 10 | Flucloxacillin | 8 μM | HL60 | 0.836 |
| 11 | Cefepime | 7 μM | PC3 | 0.836 |
| 12 | Tropine | 28 μM | PC3 | 0.834 |
| **13** | **Prochlorperazine** | **7 μM** | **HL60** | **0.825** |
| 14 | Chlorcyclizine | 12 μM | HL60 | 0.825 |
| 15 | Moracizine | 9 μM | HL60 | 0.807 |
| 16 | Lansoprazole | 11 μM | HL60 | 0.802 |
| 6006 | Cyclopenthiazide | 11 μM | PC3 | -0.8 |
| 6007 | Bromocriptine | 5 μM | HL60 | -0.8 |
| 6008 | Fluorocurarine | 12 μM | HL60 | -0.8 |
| 6009 | Mebendazole | 14 μM | HL60 | -0.801 |
| 6010 | Pronetalol | 15 μM | HL60 | -0.802 |
| 6011 | Metamizole Sodium | 12 μM | PC3 | -0.802 |
| 6012 | Profenamine | 11 μM | HL60 | -0.803 |
| 6013 | Meclozine | 9 μM | HL60 | -0.803 |
| 6014 | Ipratropium Bromide | 10 μM | HL60 | -0.803 |
| 6015 | Bemegride | 26 μM | PC3 | -0.805 |
| 6016 | Prestwick-685 | 11 μM | PC3 | -0.805 |
| 6017 | Etofylline | 18 μM | PC3 | -0.806 |
| 6018 | Luteolin | 14 μM | HL60 | -0.806 |
| 6019 | Cloperastine | 11 μM | HL60 | -0.806 |
| 6020 | BCB000040 | 10 μM | MCF7 | -0.81 |
| 6021 | 0317956-0000 | 10 μM | MCF7 | -0.811 |
| 6022 | Rimexolone | 11 μM | MCF7 | -0.812 |
| 6023 | Nicotinic Acid | 32 μM | HL60 | -0.814 |
| 6024 | Urapidil | 9 μM | HL60 | -0.815 |
| 6025 | Canavanine | 14 μM | MCF7 | -0.817 |
| 6026 | Sertaconazole | 8 μM | PC3 | -0.818 |
| 6027 | Fluspirilene | 8 μM | HL60 | -0.818 |
| 6028 | Sirolimus | 100 nM | MCF7 | -0.818 |

| | | | | |
|---|---|---|---|---|
| **6029** | **Haloperidol** | **10 µM** | **HL60** | **-0.818** |
| 6030 | Sirolimus | 100 nM | MCF7 | -0.819 |
| 6031 | Succinylsulfathiazole | 11 µM | HL60 | -0.819 |
| **6032** | **Promazine** | **12 µM** | **PC3** | **-0.821** |
| 6033 | LY-294002 | 10 µM | MCF7 | -0.821 |
| 6034 | Trolox C | 16 µM | MCF7 | -0.822 |
| 6035 | Skimmianine | 15 µM | HL60 | -0.822 |
| 6036 | Acetylsalicylic Acid | 100 µM | HL60 | -0.823 |
| 6037 | LY-294002 | 10 µM | PC3 | -0.824 |
| 6038 | Piribedil | 12 µM | HL60 | -0.824 |
| 6039 | Ethotoin | 20 µM | HL60 | -0.825 |
| 6040 | Cantharidin | 20 µM | HL60 | -0.825 |
| **6041** | **Chlorpromazine** | **1 µM** | **HL60** | **-0.826** |
| 6042 | Chloramphenicol | 12 µM | HL60 | -0.829 |
| 6043 | Zimeldine | 10 µM | PC3 | -0.834 |
| 6044 | Prazosin | 10 µM | HL60 | -0.835 |
| 6045 | Wortmannin | 10 nM | PC3 | -0.835 |
| 6046 | Tyloxapol | 4 µM | HL60 | -0.836 |
| 6047 | Cefaclor | 10 µM | HL60 | -0.837 |
| 6048 | 0175029-0000 | 1 µM | PC3 | -0.839 |
| 6049 | Pentamidine | 7 µM | HL60 | -0.839 |
| 6050 | Cefalexin | 11 µM | HL60 | -0.841 |
| 6051 | Aminohippuric Acid | 21 µM | HL60 | -0.841 |
| 6052 | Fusaric Acid | 22 µM | HL60 | -0.842 |
| 6053 | Fulvestrant | 1 µM | MCF7 | -0.842 |
| 6054 | Indapamide | 11 µM | PC3 | -0.842 |
| 6055 | Sulfaphenazole | 13 µM | HL60 | -0.843 |
| 6056 | Nilutamide | 13 µM | MCF7 | -0.844 |
| 6057 | Norfloxacin | 13 µM | PC3 | -0.846 |
| 6058 | Flunixin | 8 µM | HL60 | -0.846 |
| 6059 | Pipemidic Acid | 13 µM | HL60 | -0.848 |
| 6060 | Ethotoin | 20 µM | PC3 | -0.849 |
| 6061 | Tolbutamide | 15 µM | PC3 | -0.85 |
| 6062 | Meptazinol | 15 µM | HL60 | -0.852 |
| 6063 | Fursultiamine | 9 µM | HL60 | -0.855 |
| 6064 | Naringin | 7 µM | HL60 | -0.859 |
| 6065 | Napelline | 11 µM | HL60 | -0.859 |
| 6066 | Sirolimus | 100 nM | PC3 | -0.859 |
| 6067 | Tolazamide | 13 µM | HL60 | -0.86 |
| 6068 | Pivampicillin | 9 µM | HL60 | -0.861 |
| 6069 | Amylocaine | 15 µM | PC3 | -0.863 |
| 6070 | Tolbutamide | 15 µM | HL60 | -0.863 |
| 6071 | Quinostatin | 10 µM | MCF7 | -0.864 |
| 6072 | Sulfadimidine | 13 µM | HL60 | -0.871 |

| | | | | |
|---|---|---|---|---|
| 6073 | Etamsylate | 15 µM | HL60 | -0.872 |
| 6074 | Epirizole | 17 µM | PC3 | -0.874 |
| 6075 | Fluvoxamine | 9 µM | HL60 | -0.879 |
| 6076 | Rescinnamine | 6 µM | MCF7 | -0.88 |
| 6077 | Gossypol | 8 µM | HL60 | -0.882 |
| **6078** | **Fluphenazine** | **10 µM** | **PC3** | **-0.883** |
| 6079 | Tretinoin | 1 µM | HL60 | -0.889 |
| 6080 | Trifluoperazine | 8 µM | HL60 | -0.89 |
| 6081 | Metyrapone | 18 µM | HL60 | -0.891 |
| **6082** | **Trifluoperazine** | **10 µM** | **HL60** | **-0.895** |
| 6083 | Picrotoxinin | 14 µM | HL60 | -0.9 |
| 6084 | Terguride | 9 µM | HL60 | -0.9 |
| 6085 | Pempidine | 13 µM | MCF7 | -0.901 |
| 6086 | Zalcitabine | 19 µM | HL60 | -0.906 |
| 6087 | Labetalol | 11 µM | HL60 | -0.909 |
| 6088 | Lymecycline | 7 µM | HL60 | -0.91 |
| 6089 | Flavoxate | 9 µM | MCF7 | -0.911 |
| 6090 | Memantine | 19 µM | HL60 | -0.911 |
| 6091 | Wortmannin | 10 nM | MCF7 | -0.918 |
| 6092 | Sulfadiazine | 16 µM | PC3 | -0.921 |
| **6093** | **Sulpiride** | **12 µM** | **PC3** | **-0.941** |
| 6094 | Cefalotin | 10 µM | HL60 | -0.945 |
| 6095 | Mephentermine | 9 µM | HL60 | -0.952 |
| 6096 | Sirolimus | 100 nM | PC3 | -0.954 |
| 6097 | Naftopidil | 9 µM | HL60 | -0.958 |
| 6098 | LY-294002 | 10 µM | MCF7 | -0.968 |
| **6099** | **Thioridazine** | **10 µM** | **HL60** | **-0.974** |
| 6100 | Iopanoic Acid | 7 µM | HL60 | -1 |

**CMAP drugs from genes up- or down-regulated more than 1.5 times after tyrosol 30µM treated**

| Rank | CMAP Name | Dose | Cell | Score |
|---|---|---|---|---|
| 1 | Ketotifen | 9 µM | MCF7 | 1 |
| 2 | Ofloxacin | 11 µM | HL60 | 0.986 |
| 3 | Sulfamonomethoxine | 14 µM | MCF7 | 0.978 |
| 4 | Diethylstilbestrol | 15 µM | HL60 | 0.976 |
| 5 | Metaraminol | 9 µM | MCF7 | 0.976 |
| 6 | Homatropine | 11 µM | PC3 | 0.975 |
| 7 | Phentolamine | 12 µM | MCF7 | 0.97 |
| 8 | Naproxen | 17 µM | MCF7 | 0.967 |
| 9 | Metformin | 24 µM | HL60 | 0.964 |
| 10 | Benserazide | 10 µM | MCF7 | 0.962 |
| 11 | Merbromin | 5 µM | HL60 | 0.961 |
| 12 | Streptozocin | 15 µM | MCF7 | 0.96 |
| 13 | Bupivacaine | 12 µM | MCF7 | 0.952 |
| 14 | Enalapril | 8 µM | MCF7 | 0.948 |
| 15 | Iodixanol | 3 µM | MCF7 | 0.942 |
| 16 | Prestwick-1082 | 12 µM | MCF7 | 0.933 |
| 17 | Irinotecan | 100 µM | PC3 | 0.932 |
| 18 | Maprotiline | 13 µM | HL60 | 0.919 |
| 19 | Bumetanide | 11 µM | MCF7 | 0.919 |
| 20 | Atractyloside | 5 µM | HL60 | 0.919 |
| 21 | Articaine | 12 µM | PC3 | 0.917 |
| 22 | Thiamphenicol | 11 µM | PC3 | 0.916 |
| 23 | Timolol | 9 µM | MCF7 | 0.916 |
| 24 | Cefamandole | 8 µM | MCF7 | 0.916 |
| **25** | **Risperidone** | **10 µM** | **MCF7** | **0.915** |
| 26 | Isoxicam | 12 µM | PC3 | 0.914 |
| 27 | Pivmecillinam | 8 µM | MCF7 | 0.913 |
| 28 | Vincamine | 11 µM | HL60 | 0.913 |
| 29 | Acetohexamide | 12 µM | MCF7 | 0.906 |
| 30 | PF-00539758-00 | 10 µM | PC3 | 0.903 |
| 31 | Terconazole | 8 µM | PC3 | 0.901 |
| 32 | Alprostadil | 11 µM | MCF7 | 0.9 |
| 33 | dl-Alpha Tocopherol | 9 µM | HL60 | 0.897 |
| 34 | dl-PPMP | 2 µM | MCF7 | 0.896 |
| 35 | Podophyllotoxin | 10 µM | MCF7 | 0.895 |
| 36 | Nadolol | 13 µM | MCF7 | 0.894 |
| 37 | Vinburnine | 14 µM | PC3 | 0.894 |
| 38 | Prestwick-1103 | 20 µM | PC3 | 0.89 |
| 39 | Acemetacin | 10 µM | MCF7 | 0.889 |
| 40 | Merbromin | 5 µM | MCF7 | 0.887 |

| | | | | |
|---|---|---|---|---|
| 41 | Tacrine | 16 µM | MCF7 | 0.885 |
| 42 | Thiethylperazine | 6 µM | MCF7 | 0.884 |
| 43 | Naproxen | 17 µM | MCF7 | 0.881 |
| 44 | Adiphenine | 11 µM | PC3 | 0.88 |
| 45 | AR-A014418 | 10 µM | PC3 | 0.88 |
| 46 | Pepstatin | 6 µM | MCF7 | 0.879 |
| 47 | Metolazone | 11 µM | MCF7 | 0.878 |
| 48 | Yohimbine | 10 µM | MCF7 | 0.875 |
| 49 | Quinpirole | 16 µM | MCF7 | 0.874 |
| 50 | Pirenzepine | 9 µM | PC3 | 0.869 |
| 51 | Piperine | 14 µM | HL60 | 0.869 |
| 52 | Prestwick-1082 | 12 µM | PC3 | 0.868 |
| 53 | Isoflupredone | 10 µM | PC3 | 0.868 |
| 54 | Canadine | 12 µM | MCF7 | 0.865 |
| 55 | Estradiol | 10 nM | MCF7 | 0.863 |
| 56 | Iopamidol | 5 µM | MCF7 | 0.862 |
| 57 | Estropipate | 9 µM | MCF7 | 0.862 |
| 58 | Clebopride | 8 µM | MCF7 | 0.862 |
| 59 | Tetryzoline | 17 µM | MCF7 | 0.861 |
| **60** | **Levomepromazine** | **9 µM** | **PC3** | **0.86** |
| 61 | Calcium Folinate | 8 µM | MCF7 | 0.857 |
| 62 | Bicuculline | 11 µM | PC3 | 0.855 |
| 63 | Nifenazone | 13 µM | MCF7 | 0.854 |
| 64 | Dinoprost | 8 µM | MCF7 | 0.854 |
| 65 | Mephenesin | 22 µM | MCF7 | 0.851 |
| 66 | 5186223 | 12 µM | MCF7 | 0.85 |
| 67 | Sulfadimethoxine | 13 µM | MCF7 | 0.848 |
| 68 | Prednisolone | 11 µM | MCF7 | 0.848 |
| 69 | Trimipramine | 10 µM | HL60 | 0.847 |
| 70 | Bergenin | 12 µM | HL60 | 0.846 |
| 71 | Adiphenine | 11 µM | PC3 | 0.845 |
| 72 | Metampicillin | 10 µM | PC3 | 0.844 |
| 73 | Tacrolimus | 1 µM | MCF7 | 0.843 |
| 74 | Primaquine | 9 µM | HL60 | 0.843 |
| 75 | Adiphenine | 11 µM | MCF7 | 0.842 |
| 76 | Nitrendipine | 11 µM | MCF7 | 0.84 |
| 77 | Triamterene | 16 µM | PC3 | 0.84 |
| 78 | 0175029-0000 | 10 µM | MCF7 | 0.839 |
| 79 | Adiphenine | 11 µM | HL60 | 0.837 |
| 80 | Carisoprodol | 15 µM | HL60 | 0.834 |
| 81 | Fluvastatin | 9 µM | MCF7 | 0.834 |
| 82 | Alprostadil | 10 µM | MCF7 | 0.833 |
| 83 | PHA-00846566E | 10 µM | PC3 | 0.831 |
| 84 | Genistein | 10 µM | MCF7 | 0.831 |

| | | | | |
|---|---|---|---|---|
| 85 | Famotidine | 12 µM | PC3 | 0.829 |
| 86 | Dizocilpine | 12 µM | PC3 | 0.827 |
| 87 | Hyoscyamine | 14 µM | PC3 | 0.826 |
| 88 | Naftidrofuryl | 8 µM | MCF7 | 0.822 |
| 89 | Estradiol | 10 nM | MCF7 | 0.822 |
| 90 | Fluticasone | 8 µM | MCF7 | 0.821 |
| 91 | GLY-His-Lys | 1 µM | MCF7 | 0.821 |
| 92 | 5253409 | 17 µM | MCF7 | 0.82 |
| 93 | Pioglitazone | 10 µM | MCF7 | 0.82 |
| 94 | Levopropoxyphene | 7 µM | HL60 | 0.819 |
| 95 | Nadolol | 13 µM | HL60 | 0.819 |
| 96 | Hydroflumethiazide | 12 µM | PC3 | 0.819 |
| 97 | Fenbufen | 16 µM | HL60 | 0.818 |
| 98 | Meclocycline | 6 µM | MCF7 | 0.818 |
| 99 | Telenzepine | 9 µM | MCF7 | 0.816 |
| 100 | Letrozole | 14 µM | MCF7 | 0.816 |
| 101 | Diethylcarbamazine | 10 µM | MCF7 | 0.816 |
| 102 | Tocainide | 17 µM | MCF7 | 0.814 |
| 103 | Ciclacillin | 12 µM | PC3 | 0.813 |
| 104 | Noscapine | 10 µM | HL60 | 0.811 |
| 105 | Metronidazole | 23 µM | MCF7 | 0.811 |
| 106 | Adiphenine | 11 µM | MCF7 | 0.81 |
| 107 | Haloperidol | 10 µM | MCF7 | 0.81 |
| 108 | Aconitine | 6 µM | MCF7 | 0.809 |
| 109 | Fluticasone | 8 µM | MCF7 | 0.809 |
| 110 | Isoniazid | 29 µM | MCF7 | 0.808 |
| 111 | Monorden | 100 nM | PC3 | 0.807 |
| 112 | Kaempferol | 14 µM | HL60 | 0.805 |
| 113 | Hymecromone | 23 µM | HL60 | 0.805 |
| 114 | Amitriptyline | 13 µM | MCF7 | 0.805 |
| 115 | Beclometasone | 8 µM | PC3 | 0.802 |
| 116 | Streptomycin | 3 µM | MCF7 | 0.802 |
| 117 | Nifedipine | 12 µM | PC3 | 0.801 |
| **6043** | **Alpha-Ergocryptine** | **7 µM** | **PC3** | **-0.8** |
| 6044 | 8-Azaguanine | 26 µM | HL60 | -0.8 |
| 6045 | Rolipram | 15 µM | MCF7 | -0.801 |
| 6046 | Prilocaine | 16 µM | MCF7 | -0.802 |
| 6047 | Mephentermine | 9 µM | HL60 | -0.803 |
| **6048** | **Haloperidol** | **10 µM** | **MCF7** | **-0.807** |
| 6049 | Vorinostat | 10 µM | MCF7 | -0.809 |
| 6050 | Rottlerin | 10 µM | MCF7 | -0.811 |
| 6051 | Trichostatin A | 100 nM | MCF7 | -0.811 |
| 6052 | Trichostatin A | 100 nM | MCF7 | -0.811 |
| 6053 | Sirolimus | 100 nM | MCF7 | -0.812 |

| | | | | |
|---|---|---|---|---|
| 6054 | 8-Azaguanine | 26 µM | MCF7 | -0.813 |
| 6055 | LY-294002 | 10 µM | MCF7 | -0.813 |
| **6056** | **Thioridazine** | **10 µM** | **MCF7** | **-0.814** |
| 6057 | 5252917 | 14 µM | MCF7 | -0.815 |
| 6058 | Metergoline | 10 µM | MCF7 | -0.816 |
| 6059 | Dexverapamil | 10 µM | MCF7 | -0.817 |
| 6060 | Sirolimus | 100 nM | MCF7 | -0.826 |
| 6061 | Etodolac | 14 µM | PC3 | -0.831 |
| 6062 | Trichostatin A | 100 nM | MCF7 | -0.831 |
| 6063 | LY-294002 | 10 µM | MCF7 | -0.833 |
| 6064 | Phenazopyridine | 16 µM | MCF7 | -0.835 |
| 6065 | Sirolimus | 100 nM | MCF7 | -0.835 |
| 6066 | Vorinostat | 10 µM | MCF7 | -0.841 |
| 6067 | Valproic Acid | 50 µM | PC3 | -0.842 |
| 6068 | Ethotoin | 20 µM | MCF7 | -0.843 |
| 6069 | Clomifene | 7 µM | MCF7 | -0.843 |
| 6070 | Hydroflumethiazide | 12 µM | MCF7 | -0.844 |
| 6071 | Berberine | 11 µM | MCF7 | -0.844 |
| 6072 | Trichostatin A | 100 nM | MCF7 | -0.846 |
| 6073 | Trichostatin A | 100 nM | MCF7 | -0.849 |
| 6074 | Trichostatin A | 100 nM | MCF7 | -0.849 |
| 6075 | Pargyline | 20 µM | MCF7 | -0.85 |
| 6076 | Trichostatin A | 100 nM | MCF7 | -0.85 |
| 6077 | Trichostatin A | 100 nM | MCF7 | -0.851 |
| 6078 | Trichostatin A | 1 µM | MCF7 | -0.855 |
| 6079 | LY-294002 | 10 µM | MCF7 | -0.857 |
| 6080 | LY-294002 | 10 µM | MCF7 | -0.861 |
| 6081 | Dihydrostreptomycin | 3 µM | MCF7 | -0.863 |
| 6082 | Trichostatin A | 100 nM | MCF7 | -0.874 |
| 6083 | LY-294002 | 10 µM | MCF7 | -0.876 |
| 6084 | Wortmannin | 10 nM | MCF7 | -0.877 |
| 6085 | Sirolimus | 100 nM | MCF7 | -0.88 |
| 6086 | dl-Thiorphan | 16 µM | MCF7 | -0.881 |
| 6087 | Methylbenzethonium Chloride | 9 µM | MCF7 | -0.882 |
| 6088 | Trichostatin A | 100 nM | MCF7 | -0.887 |
| 6089 | LY-294002 | 10 µM | PC3 | -0.89 |
| 6090 | Procainamide | 15 µM | MCF7 | -0.891 |
| 6091 | Norfloxacin | 13 µM | PC3 | -0.892 |
| 6092 | Prestwick-559 | 8 µM | MCF7 | -0.895 |
| 6093 | Benzethonium Chloride | 9 µM | MCF7 | -0.899 |
| 6094 | Trichostatin A | 100 nM | MCF7 | -0.91 |
| 6095 | Trichostatin A | 100 nM | MCF7 | -0.913 |
| 6096 | Quinostatin | 10 µM | MCF7 | -0.918 |
| 6097 | Trichostatin A | 100 nM | MCF7 | -0.926 |

| | | | | |
|---|---|---|---|---|
| 6098 | Trichostatin A | 100 nM | MCF7 | -0.935 |
| 6099 | Trichostatin A | 100 nM | MCF7 | -0.944 |
| 6100 | LY-294002 | 10 µM | MCF7 | -1 |
